# Ultrafiltration Segregates Tissue Regenerative Stimuli Harboured Within and Independent of Extracellular Vesicles

**DOI:** 10.1101/2020.01.28.923037

**Authors:** TT Cooper, SE Sherman, T Dayarathna, GI Bell, Jun Ma, DM McRae, F Lagugné-Labarthet, SH Pasternak, GA Lajoie, DA Hess

**Affiliations:** Department of Physiology and Pharmacology, Western University, London, Ontario, Canada; Molecular Medicine Research Laboratories, Robarts Research Institute, London, Ontario, Canada; Don Rix Protein Identification Facility, Department of Biochemistry, Western University, London, Ontario, Canada; Department of Chemistry, Western University, London, Ontario, Canada

## Abstract

The release of extracellular vesicles (EVs) from human multipotent stromal cells (MSC) has been proposed as a mechanism by which MSC mediate regenerative functions *in vivo*. Our recent work has characterized MSC derived from human pancreatic tissues (Panc-MSC) that generated a tissue regenerative secretome. Despite these advancements, it remains unknown whether regenerative stimuli are released independent or within extracellular vesicles. Herein, this study demonstrates ultrafiltration is a simple method to enrich for EVs which can be injected in murine models of tissue regeneration. The enrichment of EVs from Panc-MSC conditioned media (CM) was validated using nanoscale flow cytometry and atomic force microscopy; in addition to the exclusive detection of classical EV-markers CD9, CD81, CD63 using label-free mass spectrometry. Additionally, we identified several pro-regenerative stimuli, such as WNT5A or ANGPT1, exclusive to EV-enriched CM. Endothelial cell tubule formation was enhanced in response to both Panc-MSC CM fractions *in vitro* yet only intramuscular injection of EV-enriched CM demonstrated vascular regenerative functions in NOD/SCID mice with unilateral hind-limb ischemia (*<p<0.05). Furthermore, both EV-depleted and EV-enriched CM reduced hyperglycemia following intrapancreatic injection in hyperglycemic mice (**p<0.01). Collectively, understanding the functional synergy between compartments of the secretome is required to advance cell-free biotherapeutics into applications of regenerative medicine.

## INTRODUCTION

Multipotent stromal cells (MSC) have been referred to as the ‘paramedics’ of the body^1, 2^; however, the translation of clinical efficacy from preclinical studies utilizing direct or systemic transplantation of MSC has not fulfilled expectations^3^. The failure of MSC to recapitulate the therapeutic effects demonstrated in pre-clinical models, has been accounted for by transient cell engraftment^4^, poor biodistribution^5–7^, and/or the loss of a pro-regenerative secretome^8–11^. Alternatively, we have recently demonstrated bone marrow (BM) MSC may be utilized as ‘biofactories’ to generate a potent serum-free cocktail capable of stimulating pancreatic islet regeneration in hyperglycemic NOD/SCID mice following intrapancreatic injection^12, 13^. The use of a cell-free biotherapeutic approach mitigated deficiencies with direct cell transplantation^14^; yet additional investigations remain necessary to establish the safety and to maximize the efficacy of this approach.

Deciphering the phenotypic segregation between human MSC and MSC-progeny, such as fibroblasts, is continuously under scrutiny^15–17^. As such, the capacity for MSC to stimulate tissue regeneration has been proposed as a defining characteristic to segregate MSC from non-therapeutic counterparts^18, 19^. We recently conducted an in-depth proteomic characterization of MSC derived from human pancreatic tissue preperations (Panc-MSC) that shared classical MSC characteristics with BM-MSC, albeit distinct phenotypic, secretory and functional characteristics were observed *in vitro*^20^. Interestingly, both MSC-types generated a secretome that supported endothelial and pancreatic cell functions *in vivo*. Ultimately, this data contributes to accumulating evidence that MSC generate a tissue regenerative secretome, regardless of tissue origin^14, 21^.

It is well established the secretome of MSC is enriched with bioactive stimuli, derived from amino acids, lipids or nucleic acids, that mediate regenerative processes within target cell populations.^14, 22^. Notably, EVs provide regenerative stimuli protection within a lipid bi-layer protected from enzymatic degradation. Thus, EVs can act locally in a paracrine manner or travel through systemic circulation to act on distant cell populations^23–25^. Classification of therapeutic EVs is complex and is primarily based on size and/or cellular-origin^26–28^. Although, it is well established therapeutic EVs are phenotypically and compositionally distinct from apoptotic bodies^29^. Exosomes (50nm-100nm), derived through the endosomal pathway, are released into the extracellular microenvironment following fusion of multivesicular bodies with the plasma membrane. Alternatively, microvesicles (100nm-1µm) are generated from blebbing of cytoplasmic compartments and localized ‘pinching’ of the plasma membrane, leading to the direct release into the surrounding microenvironment^30^. Interestingly, the luminal and membrane-bound cargo of EVs remains distinct from the composition of the parent cells^29^; thus, EV biogenesis and secretion is a highly regulated process which continues to be elucidated with technological advancements and multidisciplinary efforts^31^. As EV-based biotherapeutics are currently being tested in early human clinical trials^32^, it remains essential to further our basic understanding of EV biology while exploring diverse approaches for regenerative medicine^33^. Thus, the primary objectives of this study were 1) to provide novel insights towards proteomic cargo secreted by Panc-MSC *in vitro* and 2) assess the therapeutic potential of Panc-MSC EVs *in vivo*.

MSC-generated EVs are a potential source and delivery vehicle of therapeutic stimuli^34^, yet additional investigations remain to 1) further understand mechanisms of cellular communication driving tissue regeneration and 2) improve the production of cell-free biotherapeutics. Herein, this study employs a simple and scalable method to simultaneously enrich for and deplete EVs from conditioned media (CM) generated by MSC. The enrichment of EVs was validated by nanoscale flow cytometry, atomic force microscopy, label-free mass spectrometry, and a series of functional analyses to demonstrate the lateral transfer of luminal cargo *in vitro*. We provide novel evidence that therapeutic stimuli released from human Panc-MSC are harboured within and/or independent of EVs. Collectively, this study supports our ongoing efforts to develop novel biotherapeutics for applications of regenerative medicine to treat diabetes and its cardiovascular complications.

## METHODS

### Ultrafiltration of MSC-generated Conditioned Media

All studies were approved by the Human Research Ethics Board at the University of Western Ontario (REB 12934, 12252E). Human pancreatic islets were provided by the National Institute of Diabetes and Digestive and Kidney Diseases (NIDDK) funded Integrated Islet Distribution Program (IIDP) at the City of Hope (California, USA), NIH Grant #2EC4DK098085-02.Panc-MSC were established, cultured and used to generate conditioned media (CM), as previously described^20^. CM was subsequently processed in one of three ways **(Supplemental Figure 1)**. 1) CM was concentrated by ultrafiltration in 3kDa centrifuge filter units (Millipore) for 45 minutes at 2800g. This fraction (bulk) contained both extracellular vesicles and soluble proteins >3kDa. 2) CM was concentrated in 100kDa centrifuge filter units (Millipore) for 20 minutes at 2800 x g. This fraction (EV+) was enriched with EVs and proteins or complexes larger than 100kDa. 3) CM which passed through the 100kDa filter was centrifuged in 3kDA centrifuge filter units for 40 minutes at 2800 x g to concentrate EV-independent proteins <100KDA and >3kDa. 20mL of unprocessed CM were concentrated in 10mL batches twice to produce ∼250-300µl of bulk CM or EV-CM, whereas 120µl of EV+ CM was typically generated from 20mL of Panc-MSC CM.

### Nanoscale Flow Cytometry

Bulk, EV+, and EV-CM generated from Panc-MSC were analyzed for the number of microparticles/*μ*l. 2, 5, 10 or 20*μ*l of each concentrate was diluted to 300*μ*l with 0.22um-filtered PBS. Low-attachment 96-well plates were stored at 4°C prior to analysis at room temperature (RT). EVs were enumerated in duplicate on the Apogee A-60 micro plus nanoscale flow cytometer (nFC) with autosampler, capable of EV resolution between 150nm-1000nm^35^. 100*μ*l of diluted CM was injected and analyzed at 10.5*μ*l/min for 1 minute. The size of secreted microparticles was estimated using silica beads ranging 110nm-1300nm using properties of large-angle light scatter (LALS) and small-angle light scatter (SALS), as previously reported^35^ **(Supplemental Figure 2)**. Silica beads provide a refractive index (λ=1.42) that is closer to cells (λ=1.35-1.39) than commonly used polystyrene beads (λ= 1.59). The resolution of exosomes (<100nm) from background noise was unattainable based on the properties of the A-60 nanoscale flow cytometer during this study. In order to assess protein expression on EVs, EV- or EV+ CM was incubated with conjugated antibodies in a 1:1 ratio that was diluted 5-fold with PBS prior to incubation at overnight at 4°C. Following incubation, samples were diluted to 300µl and analyzed by nFC as described above. Parent cells were stained in parallel and analyzed by flow cytometry using a LSRII flow cytometer at London Regional Flow Cytometry Facility. Conventional flow cytometry and nFC data were analyzed using FlowJo v10.2. Antibodies and consumables are listed in **Supplemental Table 1**.

### Protein and RNA Quantification of Panc-MSC EV- and EV+ CM

CM fractions from MSC populations were analyzed for total protein content using the detergent compatible reagent to measure 660nm absorbance. The membranes of extracellular vesicles were ruptured with a urea-based lysis buffer^36^ containing ammonium bicarbonate, DTT, and 20% SDS at a 1:1 ratio followed by tip probe sonication. Lysed CM was then diluted 200-fold with an ionic detergent compatibility reagent (Thermo Fisher) and incubated for 5 minutes at RT. RNA was extracted from a normalized 50µg of EV+ or EV- CM using a RNeasy Plus Mini Kit (Qiagen), according to manufacture’s instructions. RNA was eluted in 30µl of RNAase-free ddH_2_O and stored at −80°C prior to quantification using the ND 1000 nanodrop spectrophotometer.

### Atomic Force Microscopy

EV+ and EV- CM fractions were washed twice and diluted 1:10 with 0.22µm filtered PBS. CM fractions were pipetted as 10µL microdroplets onto sterile glass coverslips and allowed to dry at RT in a sterile biological cabinet in preparation for atomic force microscopy (AFM) imaging. AFM measurements were performed using a BioScope Catalyst AFM (Bruker) equipped with NCL tips (NanoWorld) using Nanoscope software. Images were recorded in noncontact mode in air at a line rate of 1 Hz and processed using the post-acquisition software Gwyddion.

### Label-free Mass Spectrometry and Annotated Enrichment Analyses-

Methodological details of protein precipitation, digestion, liquid chromatography-tandem mass spectrometry, and proteomic data base analysis are described thoroughly in previous studies^13, 36, 37^. LC-MS/MS settings are outlined in **Supplemental Table 2**. Annotated enrichment analyses were performed using open-source Metascape^38^ (metascape.org), Enrichr^39^ (amp.pharm.mssm.edu/Enrichr/), or FunRich (funrich.org) resources.

### Cell Tracker CTMPX Labelling of Panc-MSC EVs

Panc-MSC CM was incubated for 40 minutes at RT with 1µM of Cell Tracker CTMPX, a fluorescent dye that becomes a membrane impermeable product by esterase-activity. CTMPX-labelled CM was processed to generate EV+ and EV- CM. EV- CM was used as a negative control, as this fraction is devoid of EVs capable of retaining CTMPX metabolites. CMPTX+ EVs were analyzed by nFC (see above) and visualized using oil immersion confocal imaging at 63x with 4x digital zoom (Leica TSP8).

### Human Microvascular Endothelial Cell Uptake of Panc-MSC EVs

Human Microvascular Endothelial Cells (HMVEC) were seeded at 9.40×10^3^ cells/cm^2^ in complete endothelial growth media, EGM-2 (EBM-2 + 5%FBS, IGF, bFGF, EGF, and VEGF) for 24 hours. HMVEC were washed twice with PBS and cultured in serum/growth factor-deprived EBM-2 for up to 48 hours. To show the rapid binding of EVs to HMVEC, CTMPX-labelled EV+ CM was spiked at increasing doses (15,30,45ug) into single-cell suspensions of 1.0 x 10^6^/mL HMVEC and immediately analyzed for geometric mean fluorescent intensity (MFI) via flow cytometry during a 400 second recording. Furthermore, CTMPX-labelled EVs were used to semi-quantify the rate of uptake of EVs by HMVEC using flow cytometry. CTMPX+ EV uptake was demonstrated by adding CTMPX+ EV+ or EV- CM to serum-free HMVEC cultures for up to 12 hours. MFI was measured at 0.5, 1, and 12 hours.

Confocal images were acquired at 1 or 12 hours after CFMPM-labelled HMVEC were fixed using 10% formalin and counterstained with DAPI. Alternatively, saponin-permeabilized HMVEC were stained for cytoplasmic actin filaments using phalloidin ifluor488 and counterstained with DAPI. EV internalization were visualized through the Z-plane at 0.1*μ*m increments under 63X oil immersion and 2X or 5X digital zoom (Leica TSP8). Line scan analyses were collected using Fiji software (ImageJ) and 3D volume projections were performed using LASX software (Leica).

### Assessment of Tubule Formation

In order to understand the relevance of EV-uptake in cultured HMVEC, we assessed HMVEC tubule formation under serum starved conditions in the presence and absence of bulk, EV+ or EV- CM. 1.2×10^5^ HMVEC were cultured on growth factor-reduced Geltrex (LifeTechnologies) in EBM-2 or EBM-2 supplemented with 30µg of bulk, EV+ or EV- CM generated by BM-MSC or Panc-MSC. At 24 hours, four photo-micrographs were taken per well, and complete tube formation was enumerated by manual counting in a blinded fashion using Fiji software (ImageJ).

### Femoral Artery Ligation and Intramuscular Injection

Animal procedures were approved by the Animal Care Committee at the University of Western Ontario according to guidelines of the Animal Use Protocol (2015-012).Unilateral hind limb ischemia was induced in anesthetised (100mg/kg ketamine/xylazine, maintain with 2% isoflurane, 0.8L/min) NOD/SCID mice via surgical ligation and cauterization of the femoral artery and vein (FAL), as previously described^40^. NOD/SCID mice with unilateral hind-limb ischemia (perfusion ratio <0.1), were injected 24-hours after surgery (Day 0) with a total of 80*μ*g by intramuscular (i.m) injections (20*μ*L/injection) at 3 sites in the thigh muscles and one injection in the calf muscle. To serve as a vehicle control, mice received i.m.-injections of basal AmnioMAX C100 in a similar fashion. Anesthetised NOD/SCID mice were warmed to 37°C for 5 minutes and hindlimb blood perfusion was measured using Laser Doppler perfusion imaging (LDPI) in a blinded fashion. Recovery of blood flow was quantified on 1-, 3-, 7-, 14-, and 21- and 28-days post-transplantation by comparing the perfusion ratio (ischemic/non-ischemic) of injected mice.

### Streptozotocin-induced Hyperglycemia and Intrapancreatic Injection

Animal procedures were approved by the Animal Care Committee at the University of Western Ontario according to guidelines of the Animal Use Protocol (2015-033).Pancreatic β-cell ablation and resting hyperglycemia was induced in NOD/SCID mice aged 8- to 10-week-old (Jackson Laboratories, Bar Harbor, ME, USA) by intraperitoneal injection of streptozotocin (35 mg/kg/day) for 5 consecutive days, as previously described^41^. On Day 10, hyperglycemic (15–25 mmol/l) mice were transplanted by intrapancreatic injection^41^ of basal AmnioMAX C-100 or 8µg total protein of concentrated CM. Non-fasted systemic blood glycemic levels were monitored in a blinded-fashion weekly for up to 42 days.

### Statistical Analyses

Analyses of significance were performed by paired student’s t-test or one-way ANOVA with Tukey’s multiple comparisons tests for in vitro or *in vivo* experiments. Data was exclusively compared within timepoints and between conditions. GraphPad (Prism) software was used for statistical analyses unless otherwise stated. Analyses of proteomic differences between EV+ and EV- CM were performed in Perseus software. Proteins were considered significant if enriched >2-fold using permutation false-discovery (p<0.05). Significance of enrichment annotations were determined by Metascape, FunRich or Enrichr annotated enrichment resources.

## RESULTS

### Ultrafiltration enriched for EVs generated by human Panc-MSC

We have recently demonstrated that the secretome of Panc-MSC contains vesicle-like structures which harbour cargoes including proteins, lipids, and nucleic using custom fabricated nanohole trapping and surface enhanced Raman spectrometry^22^. Ultrafiltration was used to concentrate CM; however, it remains unknown if this method could be modified to effectively segregate EVs from EV-independent cargo within the secretome of Panc-MSC. Accordingly, we validated the segregation of EV+ and EV- CM by measuring EV content using nFC analyses **(Figure 1A).** An increasing linear detection of EVs (170nm-1*μ*m) was only observed with increasing amounts of EV+ CM, whereas the number EV- CM events did not surpass background measurements of basal media **(Figure 1B).** The majority of detected EVs in EV+ CM were estimated to be smaller than 300nm relative to silica bead standards (**Figure 1C; Supplemental Figure 2**). Indeed, AFM validated the enrichment of 3-demensional vesicle structures in EV+ CM **(Figure 1D)**. The dimensionalities of detected structures, in addition to cup or dome-shaped architecture, were consistent with previous studies identifying EVs with AFM^42, 43^. Notably, we did not observe parallel vesicle structures in EV- CM. The lipid bilayer of EVs provides protection of nucleic acid cargo, including DNA, mRNA, and microRNA, from degradation and facilitates lateral transfer of cargo to recipient cell populations^34^. Accordingly, we demonstrated a ∼2-fold enrichment of RNA content within EV+ CM compared to EV- CM **(Figure 1E)**. On the other hand, the majority of the protein mass (∼70%) secreted by Panc-MSC was detected within EV- CM **(Figure 1F)**. Collectively, these results demonstrate ultrafiltration is a simple method to segregate EVs from vesicle-independent cargo secreted by Panc-MSC.

**Figure 1.**
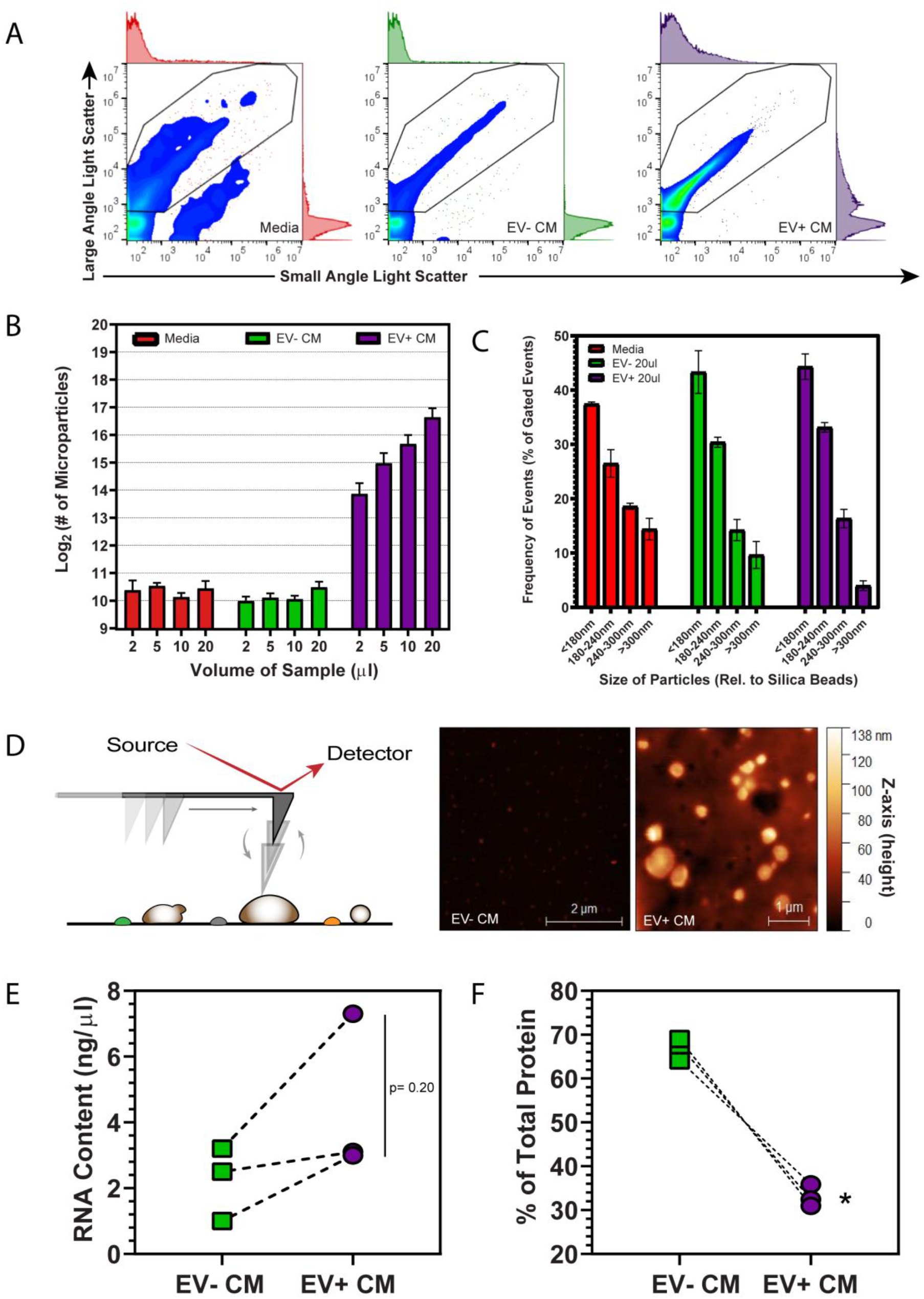
Linear detection of extracellular vesicles generated by Panc-MSC. (A) Representative nanoscale flow cytometry plots and adjunct histograms acquired using Apogee A-50, highlighted the selection of EVs generated by human Panc-MSC. Regions of EVs were gated based on large-versus small- angle light scatter intensities calibrated using silica beads ranging from 178 to 1300nm (see Supplemental Figure 2A). All samples were diluted to 300µl with 0.22µm filtered PBS and 100µl was acquired using the Apogee A-50 nanoscale flow cytometer. (B) Only EV+ CM produced an increasing linear pattern of EV detection when increasing amounts of samples were analyzed. In contrast, the total number of EVs detected in EV- CM did not exceed background detection rates, compared to concentrated media of equal volume. (C) The majority of enumerated EVs were estimated to be <300nm, relative to silica bead standards. (D) Tapping-mode atomic force microscopy was used to visualize the 3D architecture of EVs following ultrafiltration. Representative AFM recordings demonstrate EV+ CM contained vesicle structures with 3-demensional properties consistent of EVs. Parallel structures were not observed in EV- CM. (E) EV+ CM demonstrated an increasing enrichment of RNA content compared to EV- CM. (F) Alternatively, the bulk mass of protein was contained within EV- CM. (*p<0.05; n=3). Significance of analyses were determined by paired student’s t-test comparing EV- and EV+ CM.

### Classical EV-markers was exclusive to EV+ CM

EVs harbour cargo unique to the parent cell; however, less is known about which proteins are exclusively released via EVs versus EV-independent proteins during the generation of CM. We employed label-free LC-MS/MS to investigate exclusive and differential differences within the proteomic composition of EV- versus EV+ CM. Although EV- CM contained the majority of protein mass **(see Figure 1H)**, we determined the diversity of proteomic cargo within EV+ CM was significant compared to EV- CM **(Figure 2A)**. Specifically, we quantified 1458±63.02 versus 877.30±78.19 unique proteins in EV+ CM versus EV- CM, respectively **(Figure 2A).** Exclusive proteins contained 342 exclusive proteins while EV- CM contained 18 exclusive proteins that contributed to a distinct proteomic make-up **(Figure 2B-C; Supplemental Table 3, 4)**. We sought to determine if the molecular weight of segregated proteins in EV- versus EV+ CM. Interestingly, the mean molecular weight of exclusive proteins in EV- or EV+ CM was less than the 100kDa molecular weight cut off **(Figure 2D).** It was expected EV- CM would contain proteins <100kDa; however, these results led us to speculate exclusive proteins in EV+ CM that were <100kDa may be contained within EVs^44^. These speculations were supported by an enrichment of proteins associated with vesicle lumens, exosomes, or cytoplasmic ribosomes often detected in the lumen of EVs in association with nucleic acid cargo^45, 46^ **(Supplemental Figure 3A-C).** In addition, several potent regenerative cytokines (e.g. HGF, ANGPT1, Wnt5A) were exclusively detected in the EV+ fraction. Notably, classical EV-markers^46^ such as CD9, CD81 and CD63 were also exclusively detected in EV+ CM **(Figure 2E, Supplemental Figure 3A, Supplemental Table 4)**. We validated CD81 and CD63 expression on EVs and parent cells using nanoscale and conventional flow cytometry **(Figure 2F-G)**, respectively. We validated the exclusive detection of THY1/CD90 and ITGA6/CD49f and demonstrated their expression on the surface of Panc-MSC EVs and parent cells **(Figure 2H-I)**. Collectively, our mass spectrometry supports the use of ultrafiltration to enrich or deplete for EVs secreted from human MSC populations.

**Figure 2.**
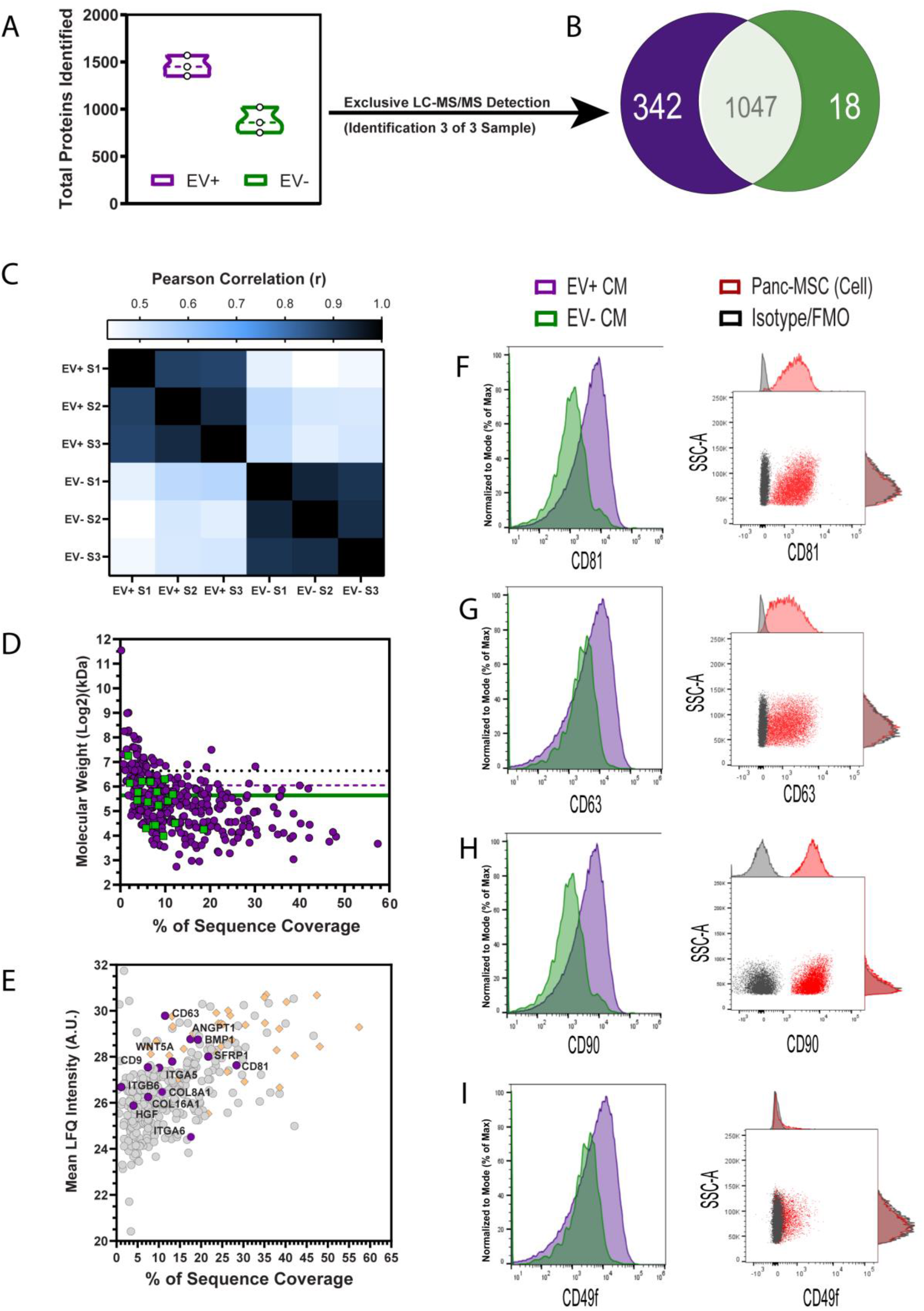
Classical EV markers (CD9, CD81, CD63) were exclusively detected in EV+ CM. Despite containing less protein by mass (see Figure 1), (A) EV+ CM contained an increased diversity of proteins compared to EV-CM determined by label-free mass spectrometry. (B) 342 or 18 of these proteins were exclusively detected within EV+ or EV- CM, respectively. Dashed line indicates median of data points. (*p<0.05; N=3). (C) Pearson correlation supports that ultrafiltration is a reproducible method to produce EV+ or EV- CM between multiple MSC donors. (D) Average molecular weight for exclusive proteins isolated in EV+ (dashed purple line) or EV- (solid green line) was compared to 100kDa ultrafiltration cutoff (black dotted line). (E) Of the 342 exclusive proteins to EV+ CM, an enrichment of ribosomal proteins (orange diamonds) was detected in addition to the exclusive identification of classical EV markers, including CD9, CD63 and CD81. Ligands of several signaling pathways, such as BMP1, ANGPT1, HGF and Wnt5a were exclusively detected in EV+ CM. Furthermore, EV+ CM exclusively contained proteins known to be expressed on the surface of Panc-MSC, such as THY1/CD90 and ITGA6/CD49f. Accordingly, we validated the detection of (F) CD81, (G) CD63, (H) CD90 and (I) CD49f on both the external membrane of EVs (left) and parent cells (right) using nanoscale for conventional flow cytometry, respectively.

### Human endothelial cells uptake CTMPX-labelled EVs generated by Panc-MSC

The exclusive fraction of EV+ CM was highly enriched with proteins associated with extracellular matrix organization and cell surface interactions at vascular wall **(Supplemental Figure 3D-F)**. Several of these proteins identified are crucial mediators of cell-cell and/or cell-matrix interactions during regenerative processes, such as ANGPT1, COL8A1 or ITGA6. EVs express integrins and/or integrin ligands^46^ to facilitate the docking of EVs to the exterior surface of cell populations expressing complementary machinery^47^, exemplified by the capacity of bone-marrow MSC to stimulate regeneration within distant renal tissues^48^. In an effort to visualize uptake of EVs within endothelial cells, labelling of EVs was accomplished with Cell Tracker^TM^ CMTPX after collection of cell-free CM and prior to ultrafiltration **(Supplemental Figure 4A)**. Accordingly, NFC validated the increased frequency and linear detection of CMTPX+ EVs within EV+ CM, compared to EV- CM **(Supplemental Figure 4B-F).**

Next, we investigated whether EV+ CM could transfer luminal cargo to human endothelial cells *in vitro*. Specifically, we sought to measure the lateral transfer of CMPTX using conventional techniques, such as flow cytometry or high-resolution confocal imaging **(Figure 3A-B)**. EV-uptake can occur through several mechanisms, such as phagocytosis^49^, macropinocytosis^50^ and clathrin- or caveolae-mediated endocytic processes^51, 52^. In order to crudely illustrate these mechanisms, HMVEC in single-cell suspension were exposed to increasing doses of CMTPX-labelled EV+ CM and were immediately assessed CMPTX accumulation over time using flow cytometry. Accumulation of CMTPX fluorescence within suspended HMVEC occurred within 400 milliseconds and increased MFI in a dose-dependent fashion **(Figure 3C)**. Furthermore, sustained uptake of CMTPX+ EVs within CMFPR-labelled HMVEC cultured with CMTPX-labelled EV+ CM was observed up to 12 hours using both flow cytometry **(Figure 3D-E)** or confocal microscopy **(Figure 3F)**. Notably, CMTPX accumulation was not observed within HMVEC supplemented with EV- CM.

**Figure 3.**
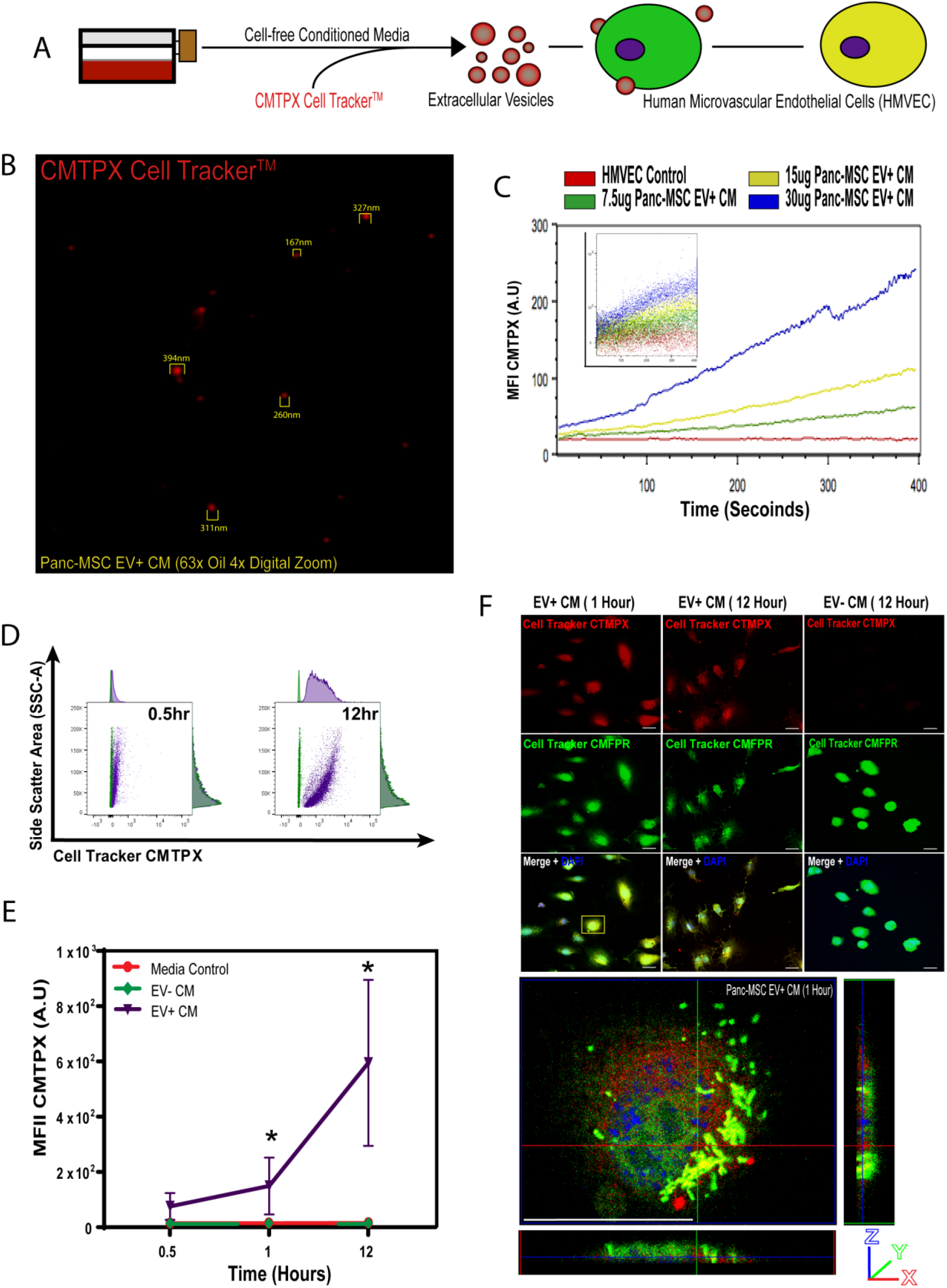
Human microvascular endothelial cells uptake CMTPX-labelled Panc-MSC EVs. (A) Schematic highlighted the cell-independent labeling of Panc-MSC EVs using CMTPX Cell Tracker^TM^ in order to assess uptake in cultured human microvascular endothelial cells labelled with CMFPR Cell Tracker^TM^. (B) Representative oil immersion confocal photomicrograph of Panc- MSC EV+ CM highlight the detection of CMTPX+ EVs. Notably, diameters of CMTPX+ EVs aligned with previous nanoscale flow cytometry analysis (see Figure 1). (C) Single-cell suspensions of HMVEC were exposed to increasing amounts of CTMPX-labelled EV+ CM and analyzed immediately for CMTPX uptake/accumulation using conventional flow cytometry up to 400ms. Specifically, the geometric mean fluorescence intensity (MFI) was used to semi-quantify CMPTX uptake by HMVEC. (C) Representative flow cytometry plots demonstrate that only the EV+ CM fraction were able to increase CTMPX fluorescence in adherent HMVEC. Accordingly, (C) CMTPX-labeled EVs were detected within adherent CMFPR+ HMVEC at 1 and 12 hours, whereas uptake of CMTPX accumulation was not observed in CMFPR+ HMVEC cultured with EV- CM. Data represented as Mean ± SEM (*p<0.05; n=3). Significance of analyses were performed by one-way ANOVA and were assessed exclusively at matched timepoints.

To confirm active uptake of EVs rather than passive transfer of CMPTX dye, we visualized EVs at the z-plane of the plasma membrane and perinuclear regions of the cytoplasm of HMVEC using compartment specific dyes, such as phalloidin to mark cytoplasmic actin **(Figure 4A-D)**. Specifically, CMTPX+ EVs were detected at the z-plane of actin filaments after five minutes of EV+ CM exposure on serum-starved HMVEC, based online segment intensity profiles **(Figure 4B, D)** and 3D volume projections **(Figure 4E).** These results suggest endothelial cells are component to the uptake of Panc-MSC EVs *in vitro*.

**Figure 4.**
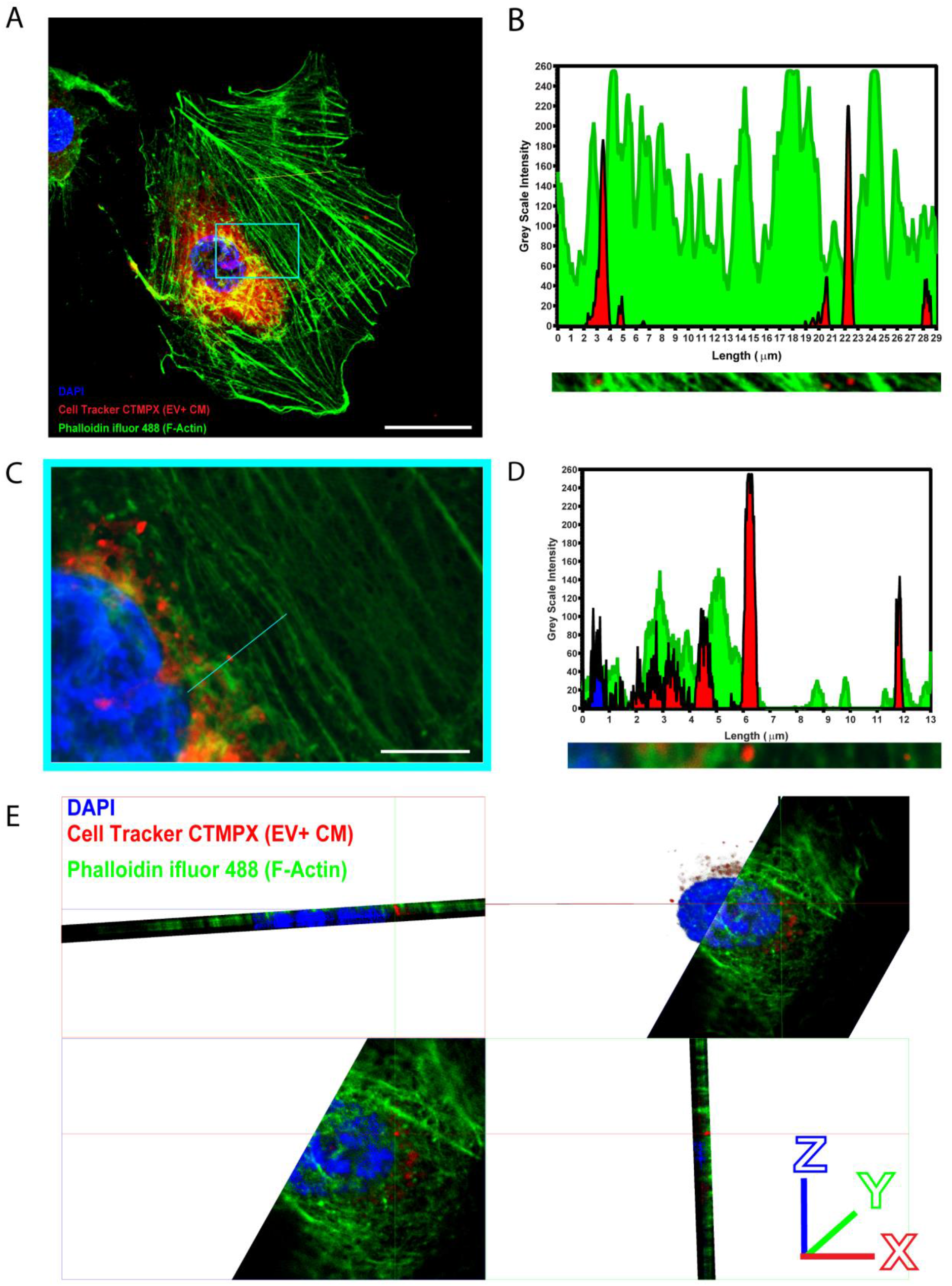
Detection of Panc-MSC EVs within the cytoplasm of human endothelial cells. Representative confocal photomicrograph of adherent HMVEC cultured with CMPTX-labelled EV+ CM for 5 minutes. CMTPX+ EVs were observed at the plane of cytosolic actin, as determined by the overlap of phalloidin iFluor ^TM^ 488 fluorescence with CMPTX+ fluorescence using line intensity profiles (yellow line) at 63x. (C) CMTPX+ EVs were detected within the perinuclear region of HMVEC (blue box). (D) Additional line intensity profile analysis at 63x plus 5x digital zoom identified CMTPX fluorescence demonstrated diameters consistent with EVs and often associated or overlapped with actin filaments. Evidence of EV accumulation within the nucleus of adherent HMVEC was not observed. (E) 3-dimensional volume projections demonstrate sustained accumulation of EVs within the cytoplasm of HMVEC after 45 minutes of culture with CMPTX-labelled EV+ CM. Scale bars = (A) 25 µm or (C) 10µm.

### The secretome of Panc-MSC demonstrated vascular and tissue regenerative properties

High-throughput *in vitro* assays provide a platform to screen for therapeutic pharmacological^53^ or biological agents^54, 55^ prior to cumbersome pre-clinical *in vivo* models. We performed endothelial tubule formation, under serum starvation, as a functional readout to assesses whether EV+ or EV- CM elicit provascular functions in a suboptimal microenvironment. At 24 hours, HMVEC formed comparable number of tubules when exposed to EV-, EV+, or bulk CM. Notably, all three conditions increased the number of tubules formed compared to vehicle controls. **(Figure 5 A-B)**. These results led us to speculate that EV-dependent and independent bioactive cargo may mediate functional changes in target cell populations of regenerative medicine (i.e. endothelial cells for ischemic conditions). We have recently demonstrated the injection of MSC-generated CM facilitates tissue regeneration *in vivo* and by-passes common limitations encountered with direct cell transplantation^12^. Therefore, we first sought to investigate the vascular regenerative potential of Panc-MSC CM following intramuscular injection into NOD/SCID mice with unilateral hindlimb ischemia **(Figure 5C).** Compared to vehicle control, bulk and EV+ CM significantly increased blood perfusion over 28 days **(Figure 5D-E)**, as determined by area under the curve (AUC) of hindlimb perfusion ratios (*p<0.05; **Figure 5F**). Mice that received i.m. injections of EV- CM demonstrated an increasing trend of blood perfusion recovery, however AUC measurements were statistically comparable to vehicle control. Collectively, these results support our proteomic analyses and provide foundational evidence that the secretome of Panc-MSC contains vascular regenerative stimuli harboured within EVs.

**Figure 5.**
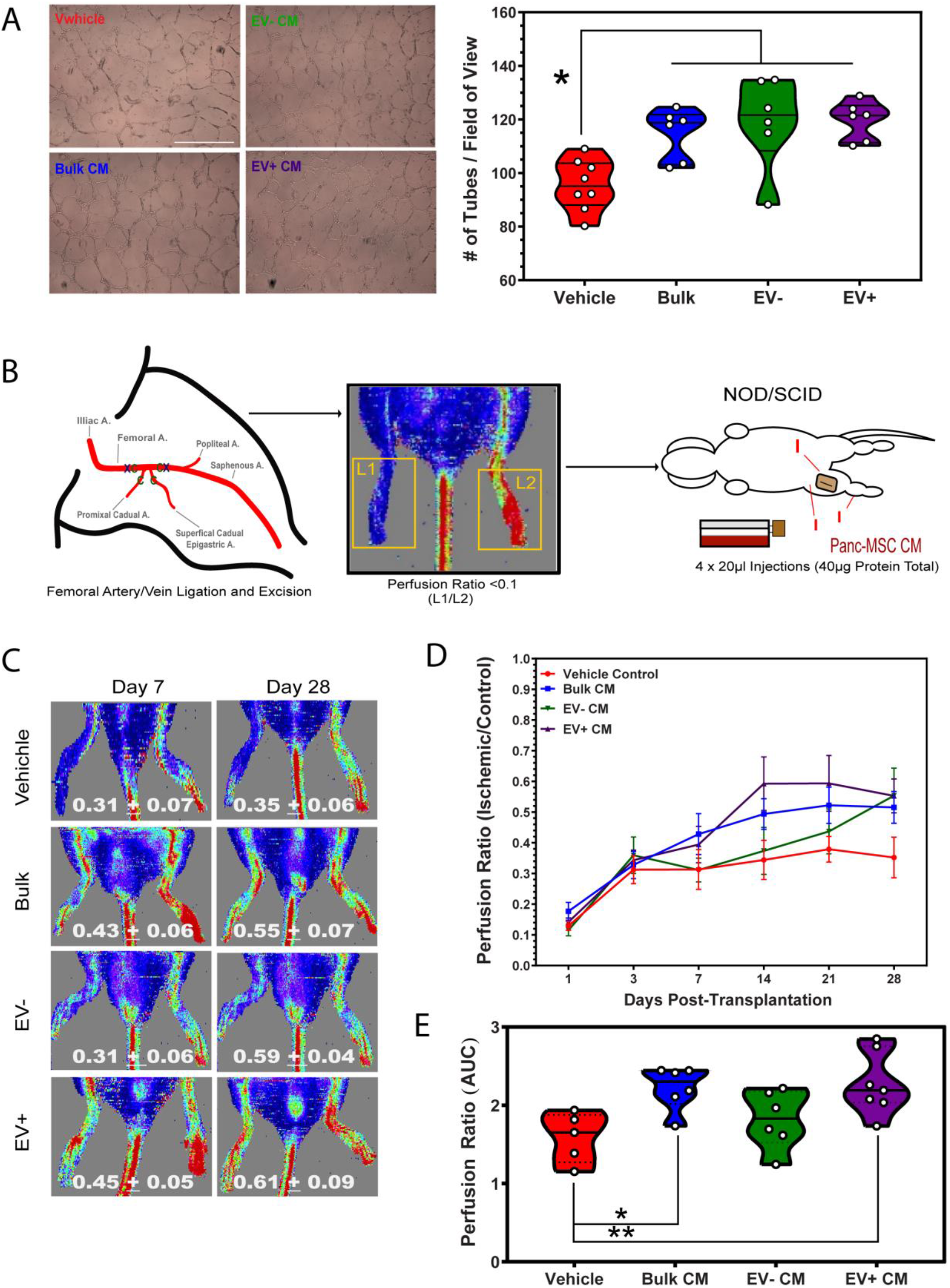
The secretome of Panc-MSC CM demonstrate vascular regenerative functions in a microenvironment of injury and acute tissue ischemia. (A) Representative photomicrographs of tubule formation by HMVEC exposed to bulk, EV-, or EV+ CM under serum-starved conditions for 24 hours. (B) Compared to HMVEC supplemented with vehicle control, basal AmnioMAX, HMVEC tubule formation was enhanced when supplemented with bulk, EV- and EV+ CM generated by Panc-MSC. Scale bar = 100µm (C) Femoral artery/vein ligation (x) and cauterization (c) was performed to induce unilateral hindlimb ischemia in NOD/SCID mice. Perfusion ratios were quantified by compared Laser Doppler perfusion imaging intensities between ischemic (L1) and non-ischemic (L2) limbs. Mice with unilateral ischemia (perfusion ratios <0.1) received intramuscular injections (I) of Panc-MSC bulk, EV-, or EV+ CM. To serve as a vehicle control, equivalent volumes of basal AmnioMAX were injected in a similar fashion. (D) Representative LDPI recordings demonstrate the enhanced recovery of (E) blood perfusion up to 28 days following intramuscular injection of bulk or EV+ CM, as determined by (F) area under the curve (AUC). EV- CM demonstrated an increasing trend, although was statistically comparable to vehicle control. Data represented as Mean ± SEM (*p<0.05; N=5). Statistical analyses were determined by one-way ANOVA with post-hoc tukey’s t-test.

Regeneration of vasculature and recovery of blood perfusion in the FAL-induced hindlimb ischemia model primarily reflects the capacity to enhance endogenous tissue regeneration. Due to the pancreatic source of Panc-MSC and associated with islets of Langerhans, we sought to assess the islet regenerative properties of EV- or EV+ CM in a more complex murine model in which endogenous regeneration has reported to be extremely limited or absent. STZ-induced β-cell ablation generates a model of chronic hyperglycemia, which demonstrates extremely limited regeneration of ablated endogenous β-cells^41, 56^. Similar to use the LDPI in the FAL model, temporal monitoring of resting blood glucoses can be utilized as a surrogate to assess the regeneration of insulin-producing β-cells **(Figure 6A)**. Notably, a single injection of bulk, EV+, or EV- CM could reduce and stabilize non-fasted resting blood glucose up 32 days post-injection, compared to injection of vehicle control **(Figure 6B-D)**, as determined by AUC. Further studies are necessary to elucidate the mechanisms of the vascular or pancreatic tissue regeneration following injection of Panc-MSC CM. Regardless, we provide novel evidence that EVs generated by Panc-MSC may be used as a biotherapeutic agents for applications of regenerative medicine.

**Figure 6.**
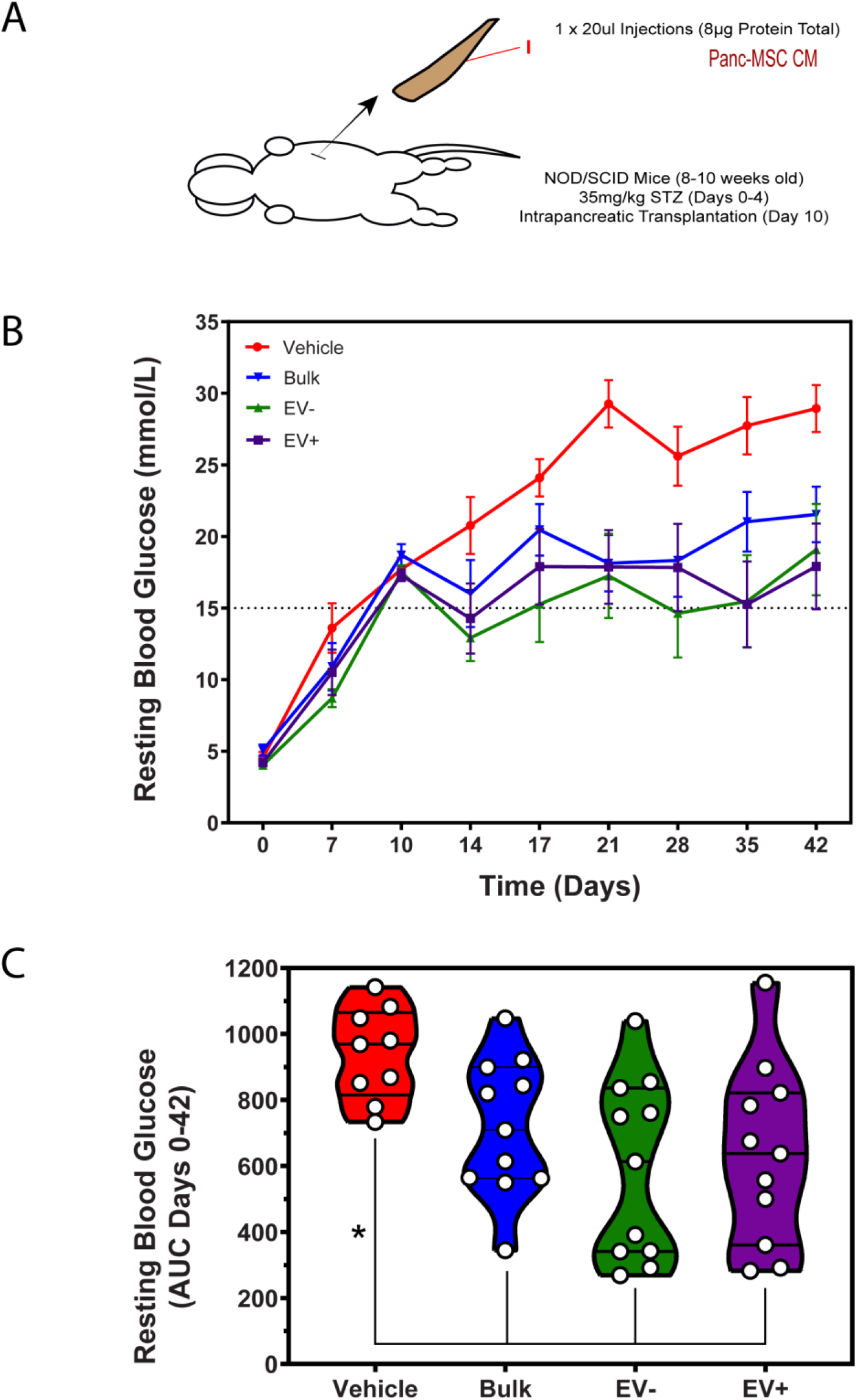
Intrapancreatic injection of Panc-MSC EV+ or EV- CM stabilizes resting blood glucose levels in streptozotocin-induced hyperglycemic mice. (A) Targeted pancreatic β-cell ablation induced in NOD/SCID mice (8-10 weeks old) via intraperitoneal injection of streptozotocin (35 mg/kg/day) between Day 0-4. On Day 10, hyperglycemic (non-fasted resting blood glucose measurements between 15–25 mmol/l) mice were transplanted in a blinded-fashion by intrapancreatic injection of AmnioMAX C-100 media or concentrated bulk EV- or EV+ CM containing ∼8µg total protein. (B) Resting blood glucoses were measured periodically up to Day 42 by tail vein puncture. (C) Intrapancreatic injection of bulk, EV-, or EV+ CM generated by Panc- MSC significantly reduced and stabilized hyperglycemia, as determined by area under the curve (AUC). Data represented as Mean ± SEM (N=3). Analyses of significance (*p<0.05) were determined by one-way ANOVA with post-hoc Tukey’s t-test

### The secretome of Panc-MSC was enriched with tissue regenerative proteins

A recent upsurge of approaches within regenerative medicine have sought to utilize the secretome of MSC as means to transfer a complex mixture of bioactive cargo to diseased or damaged tissues^34^. Considering *in vitro* tubule formation and *in vivo* tissue regeneration was supported by both EV- and EV+ CM, we hypothesized that therapeutic stimuli would be enriched within both CM fractions. 743 of 1047 proteins were detected in 2 of 3 EV+ and EV- CM samples **(Figure 7A; Supplemental Figure 5A),** albeit a distinguishable fingerprint between either fraction persisted **(Figure 7B; Supplemental Figure 5B)**. In order to perform statistical comparisons^57^, missing value imputation was performed on common proteins without compromising the integrity of the data **(Figure 7C).** Permutation-based false discovery rate (p>0.05) determined 201 proteins were significantly enriched (≥2-fold) for in the EV+ CM fraction, whereas 153 proteins were significantly enriched for in the EV- CM fraction **(Figure 7D; Supplemental Table 5, 6)**. The average molecular weight of significantly enriched proteins was ∼85kDa in ∼60kDa in EV+ CM and EV-CM, respectively **(Figure 7E)**. Supporting our previous results, EV+ CM was statistically enriched for protein associated with vesicle lumen, including COL6A2, FN1, EGFR and TFGB1 **(Figure 7F)**. Interestingly, EV- CM was also enriched for luminal proteins, such as SERPINB6 and Tissue Factor/TF **(Figure 7F)**. It is remains unclear whether these results suggest that select proteins can be secreted within or independent of EVs or if the overlap of detected proteins may be an artifact of ultracentrifugation and/or EV rupture.

**Figure 7.**
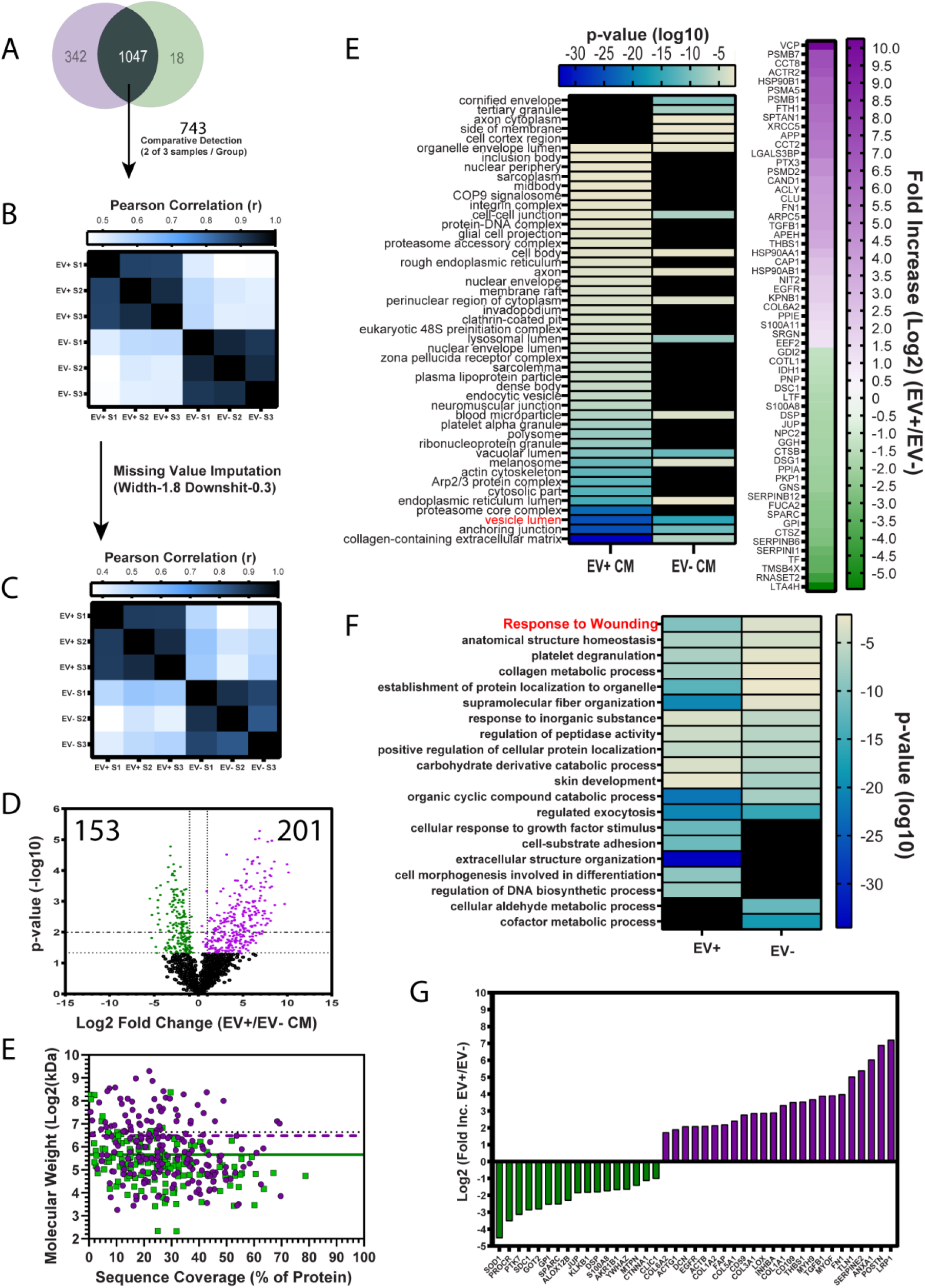
EV- and EV+ CM demonstrate distinct proteomic signatures associated with wound response and regenerative processes. Focusing on proteins detected in both EV- and EV+, (A) 743 out of 1047 proteins were identified in 2 of 3 donors of EV- and EV+ CM. Despite detection in both EV- or EV+ CM, (B) a distinct composition of these proteins existed. (C) This distinction was undisturbed by missing value imputation. (D) Specifically, 153 and 201 proteins were >2-fold enriched in either EV- and EV+ CM (p<0.05). Notably, proteins enriched within EV+ CM (purple dashed line) were slightly larger than EV- CM (green solid line), however the mean molecular weight of either was below 100kDa (black dotted line). (E) GO Cellular Compartment annotation analysis determined proteins enriched within both EV+ and EV- CM were significantly associated with the vesicle lumen (right). (F) GO Biological Processes annotation analysis determined EV+ or EV- CM were significantly enriched with proteins that are associated with (G) responses to wounding. Black boxes in (E) and (F) represent annotations which were not significant to either EV- or EV+ CM.

MSC are a fibroblastic cell population which supports homeostasis and facilitate tissue regeneration through the modulation of the immune system, promotion of angiogenesis and remodeling of the tissue microenvironment^1, 2^. With these properties in mind, we speculated serum-starvation in the microenvironment of 2D culture MSC likely mimics acute tissue injury and leads to a “wounding response” from cultured MSC^58–60^. We suspected the secretome of Panc-MSC would be enriched with tissue regenerative and matrix modifying proteins, possibly in an effort to establish homeostasis within the harsh microenvironment of culture^59^. Indeed, enriched proteins within EV- and EV+ CM are associated with responses to wounding, epithelial to mesenchymal transitions or complementary mechanisms of tissue regeneration **(Figure 7G; Supplemental Figure 5C)**. For example, we identified several known mediators of tissue regeneration and angiogenesis significantly enriched within EV+ CM, including Periostin, TGF- B1, EGFR an FN1 **(Figure 7H)**. Unexpectedly, EV- CM was significantly enriched membrane proteins, including PROCR and PTK7 **(Figure 7H)**. Nonetheless, 389 proteins were comparatively detected between EV- an EV+ CM **(Supplemental Table 7)** and were significantly associated with epithelial to mesenchymal transitions, targets of Myc signaling and angiogenesis, (**Supplemental Figure 5D).** Collectively, our early speculation that MSC generate a therapeutic secretome in response to a “wounding-like” microenvironment^61^ of 2D culture is supported by our proteomic, *in vitro* and *in vivo* analyses. Regardless, we demonstrate this response may be exploited and MSC may serve as “biofactories” to generate cell-free biotherapeutics for applications of regenerative medicine.

## DISCUSSION

This study characterized the secretome generated by Panc-MSC and determined tissue regenerative stimuli secreted by Panc-MSC were harboured within or independent of EVs. Specifically, ultrafiltration was used to simultaneously concentrate and segregate the secretome of Panc-MSC into injectable EV-enriched or EV-depleted fractions. This method was validated by nFC, AFM, and a series of *in vitro* analyses to demonstrate the lateral transfer of cargo via EVs using confocal microscopy and conventional flow cytometric analyses. In addition to augmenting endothelial tubule formation *in vitro*, direct injection of the EV-enriched or -depleted CM enhanced the recovery of blood perfusion in mice with FAL-induced hindlimb limb ischemia. Furthermore, the intrapancreatic injection of EV- or EV+ CM reduced and stabilized resting blood glucose in STZ-treated hyperglycemic mice. Overall, we provide a simple workflow to characterize therapeutic EVs generated by MSC and investigate potential applications of MSC-generated biotherapeutics.

Witwer *et al.* recently highlighted the collective focus of ISCT and ISEV to further improve the guidelines for classifying MSC and MSC-generated EVs designed for therapeutic applications^26^, respectively. Indeed, the microenvironment of culture^61^ and/or tissue source of origin^62–65^ can lead to dynamic phenotypes and functional characteristics demonstrated by MSC *in vitro*^58, 66^. It is also becoming evident that serial passaging and unwanted differentiation of MSC can lead to a loss of therapeutic activity^14, 19, 67^, such as the acquisition of a senescent secretome^68^. Thus, it appears the secretome of MSC may provide an additional layer of classification which may ultimately segregate therapeutic MSC populations from differentiated progeny^19^ or senescent counterparts^13^. The therapeutic activity of human MSC has recently been attributed to the release of EVs^26, 34^, specifically exosomes, amongst additional mechanisms of lateral transfer^69^. Despite many of these studies demonstrating a clear role for EVs towards the therapeutic functions of stem and progenitor cells, it remains unclear if bioactive stimuli are exclusive to EVs^44^.

EVs may be purified by ultracentrifugation^70^, density/size-based fractionation^71, 72^, or proprietary commercial separation kits^73^; however, each method may require uncommon reagents and/or equipment and may not preserve EV-independent components of the secretome. Alternatively, ultrafiltration appears to be a simple and scalable method to enrich EVs from MSC CM in order to screen for therapeutic function(s) *in vitro* or *in vivo*. The caveat to this method is the unavoidable co-purification of proteins or protein complexes larger that 100kDa, in addition to the disruption of EVs within pores of the filter unit. Nonetheless, ultrafiltration allowed us to demonstrate, using *in vitro* and *in vivo* models, that the transfer of therapeutic stimuli within the secretome of Panc-MSC is likely not exclusive to EV biogenesis and secretion.

The attractiveness of EVs for therapeutic applications is in part due to the expression of surface proteins that may select for and mediate uptake in recipient cells^26, 74^. The surface of EVs often comprised of proteins co-expressed on exterior and interior plasma membrane of parental cells, thus these proteins will function to mediate interactions with the extracellular matrix, recipient cells or with parental cell in an autocrine-like manner^74^. Several integrins are commonly expressed on MSC^20^ and are known machinery which facilitate EV docking and uptake^74–77^. EVs generated by Panc-MSC expressed several MSC-associated surface proteins, including THY1/CD90 and integrin ITGA6/CD49f; in addition, to classical EV-markers CD81 and CD63. Accordingly, EVs generated by Panc-MSC accumulated at the plasma membrane and within the cytoplasm of human endothelial cells, which translated to improved endothelial functions *in vitro* and *in vivo*. Future studies will need to explore the individual contribution of EV-bound proteins towards uptake and/or functional response(s) observed in recipient cell populations, such as endothelial or pancreatic cells. Albeit, our results suggest effectors independent of EVs also activate tissue regenerative mechanisms *in vivo.* This paradigm has been demonstrated in previous, such as EV-independent induction of tumor progression^78^ and Argonaut 2 complex shuttling of microRNAs in human plasma^79^.

The lipid bilayer of EVs provides a biological barrier to protect cargo from enzymatic degradation within the extracellular microenvironment of tissue or blood plasma. In addition to proteomic stimuli, nucleic acids may also mediate regeneration in similar models^34^. Losordo and colleagues demonstrated human CD34+ hematopoietic progenitors generated exosomes *in vitro* that transferred miR-126-3p to reduce SPRED1 mRNA in murine endothelial cells *in vivo*, in return enhancing hindlimb revascularization and recovery from ischemia^80^. The secretome of MSC is also known to influence the infiltration hematopoietic cells following tissue injury, including the promotion of a pro-regenerative M2 macrophage differentiation via the transfer of nucleic acid cargo^81–83^. Thus, future studies will need to determine whether cargo enriched in EV+ CM can influence the phenotypic and tissue regenerative properties of infiltrating hematopoietic, stromal, or progenitor cells. At this time, we cannot conclude whether the therapeutic stimuli contained within the secretome of Panc-MSC is exclusive to proteomic or nucleic acid cargo; albeit, bioactive lipids^84^ or metabolites^85^ may also contribute to the therapeutic functions of EVs. As the field of cell-free biotherapeutic advances, we envision multiplatform analyses to standardize proteomic, nucleic acid and lipid cargo within and independent of EVs in order to fully understand the orchestration of complex tissue regeneration induced by cell-free biotherapeutics.

A collection of studies has demonstrated the potential of utilizing MSC as EV-generating biofactories and engineering the secretome of MSC to meet therapeutic needs^37, 41, 67^. Our previous studies have supported this progression of thought and we envision the creation of a cell-free biotherapeutics ‘tailored’ to combat a given pathology. For instance, several stimuli with known pro-regenerative functions were detected in MSC CM and/or EV+ CM fraction. Wnt5a signals through both the canonical and non-canonical signaling pathways to activate pro-angiogenic pathways *in vivo* through endothelial cells^86^ or through the M2-polarization of infiltrating monocytes^86, 87^. Likewise, VEGF-A, ANGPT1 and POSTN have also demonstrated pro-regenerative function in previous studies^88–90^. We have recently demonstrated supplementation of the GSK3 kinase inhibitor (CHIR99021) to BM-MSC can enhance tissue regenerative functions of injected CM^12^; whereas, other studies have demonstrated 3D bioreactor culture^67^ or hypoxic priming^91^ may improve the tissue regenerative functions of MSC. Nonetheless, applications of cell-free biotherapeutics would benefit from exploring the directed modulation of molecular targets in recipient cells. For example, a recent clinical trial (NCT03608631) has engineered BM-MSC to produce EVs with shRNA against pancreatic cancer with KRAS^G12D^ mutations^32^. In this study, CD47+ EVs were resistant to destruction by innate immunity and selectively internalized by pancreatic cancer cells *in vivo;* in return, improving overall survival rates. Collectively, novel biotherapeutic strategies will need to consider the therapeutic cargo harboured within EVs, in addition to the biological targets and mechanisms activated in recipient cells.

### Conclusions

EV biogenesis is an efficient mechanism of cellular communication integrating multiple ligands, receptors and regulators of cellular machinery (e.g. mitochondria, ribosomes, and MSC-mRNA) that can be transferred to distant cell populations to induce biological functions^92^. Recently, the field of regenerative medicine has begun to exploit the therapeutic potential of EVs, however it remains unclear 1) which cell populations should be used to generate EVs and 2) if therapeutic stimuli are exclusive to EVs. We have addressed the latter in this study, as our results suggest tissue regenerative stimuli may be independent of EVs. Regardless, we provide novel evidence that alternative MSC populations, such as Panc-MSC, may be used to generate cell-free biotherapeutics for applications of regenerative medicine. The collection of results obtained within this study further exemplifies the importance of retaining and analyzing the complete secretome of MSC during pre-clinical studies of therapeutic EVs.

## Supporting information

Supplemental Table 1 Antibody List

Supplemental Table 2 Mass Spec Settings

Supplemental Tables 3-7 Proteomic Lists

## Acknowledgements

This work was supported by an operating grant from the Canadian Institute of Health Research (CIHR) (MOP# 378189) and Juvenile Diabetes Research Foundation (2-SRA-2015-60-Q-R) awarded to DAH, in addition to NSERC Discovery grant and Canadian Foundation for Innovation grant awarded to GAL.

---

**Supplemental Figure 1.**
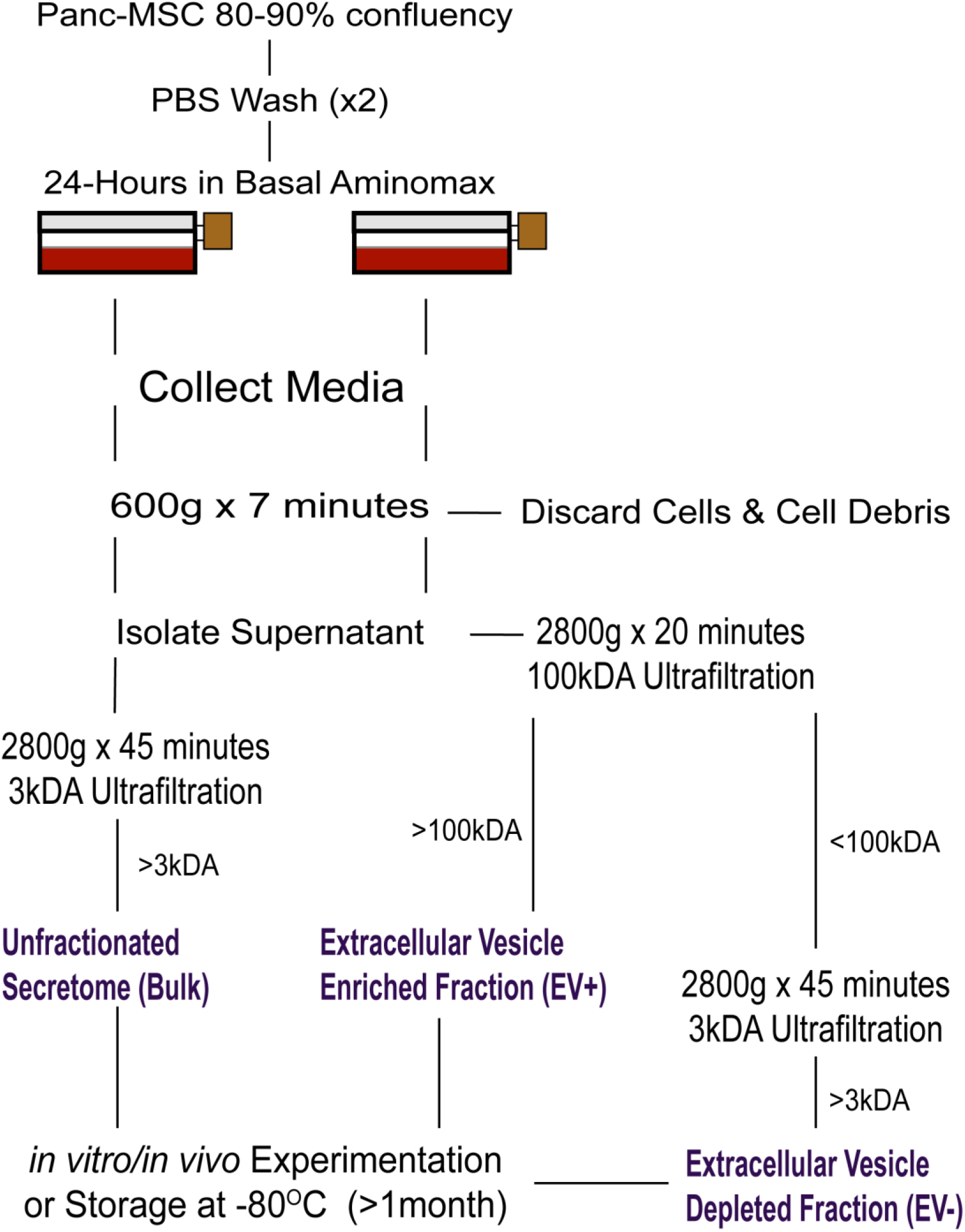
Segregation of EV+ and EV- CM using Ultrafiltration.

**Supplemental Figure 2.**
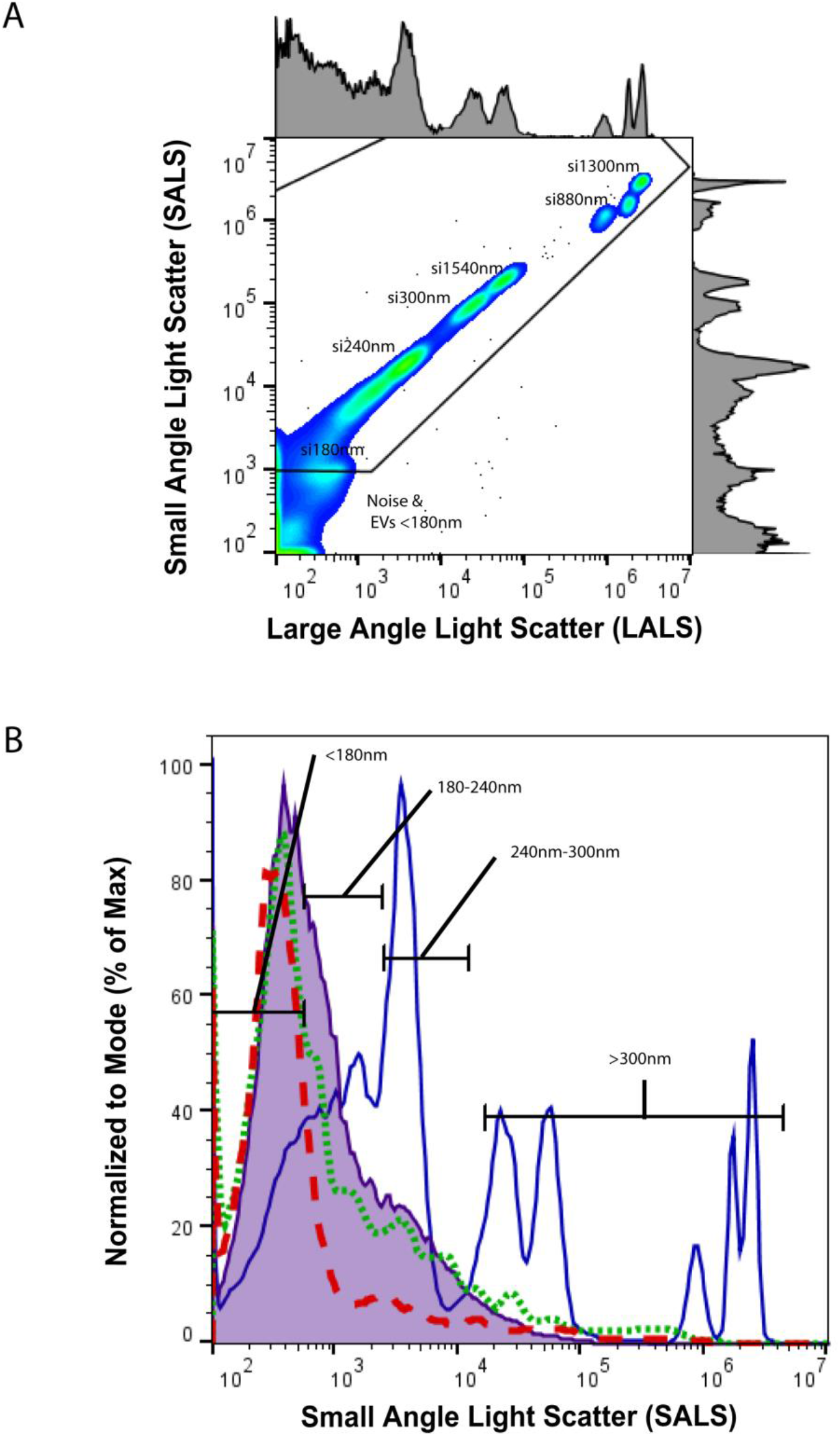
Estimation of EV size based on Silica Beads. (A) Representative histogram of silica calibration beads of various sizes. Estimated size ranges were determined using small angle light scatter properties. (B) Representative histogram demonstrating the gating strategy for EVs determine to be <180nm, 180-240nm, 240-300nm, >300nm in media only (red dashed line), EV- CM (green dotted line) or EV+ CM (purple solid) samples relative to silica beads (blue solid line) ranging from 180nm to 1300nm. Abbreviations: si = silica beads)

**Supplemental Figure 3.**
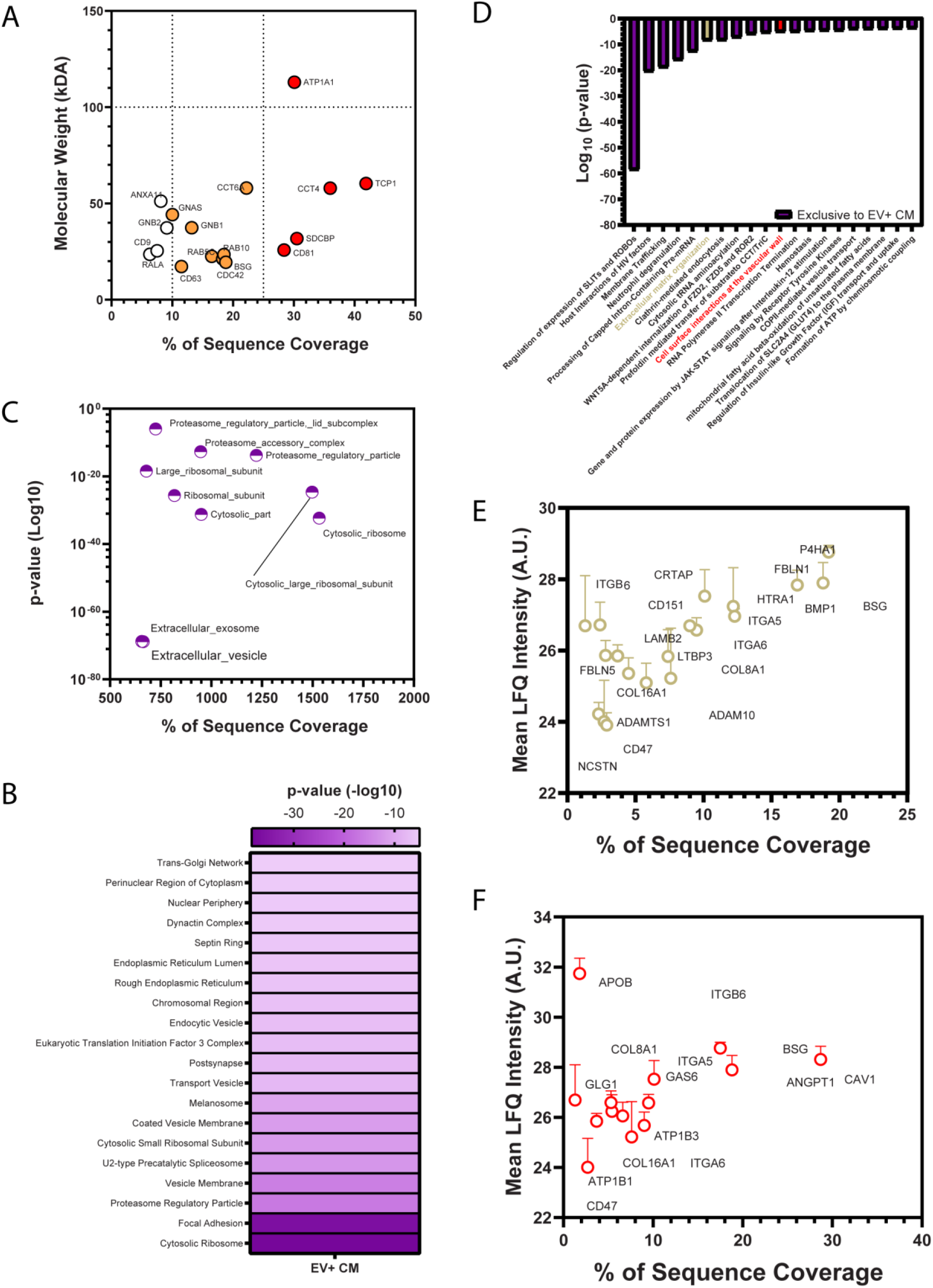
Pathway enrichment analysis of proteins exclusive to EV+ CM. Exclusive proteins detected in EV+ CM were compared against several enrichment annotation databases, including (A) Exocarta, (B) JESEN Compartments or (C) GO Cellular Components/GOCC. A significant enrichment of known exosome proteins was identified by both or Exocarta or JENSEN Compartments enrichment analyses. These included transmembrane tetraspanins embedded in the membrane of EVS, such as CD63 and CD81. In addition, proteins associated with vesicle membrane, focal adhesion or cytosolic ribosomes were significantly identified against the GOCC annotation database. Nonetheless, exclusive proteins within EV+ CM were significantly associated with several Reactome annotations, such as (E) extracellular matrix organization or (F) cellular interactions at the vascular wall. Data represented as Mean ± SD (n=3).

**Supplemental Figure 4.**
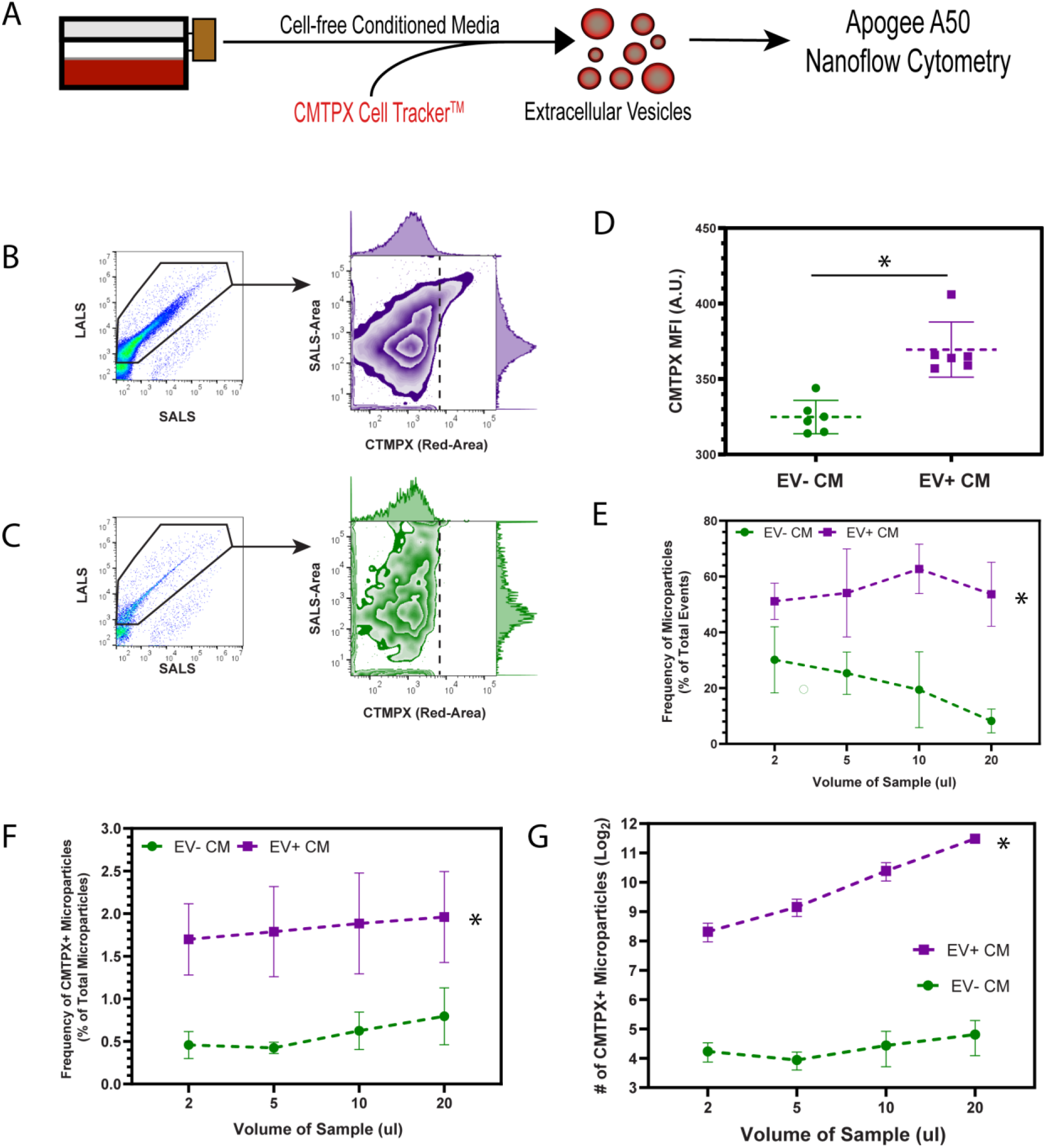
Validation of CMTPX-labelling of EVs using nanoscale flow cytometry. (A) Panc-MSC CM was stained with Cell Tracker CTMPX, allowing for the cell-independent labelling of lipid bilayer structures prior to ultrafiltration. EVs retaining CMTPX dye were detected using nanoscale flow cytometry on the Apogee A-50. Representative nanoscale flow cytometry plots of (B) EV+ or (C) EV- CM demonstrating an increase of (D) geometric mean fluorescent intensity (MFI) across gated EVs. (E) The frequency of total EVs and CMTPX+ EVs was significantly increased within EV+ CM, compared to EV- CM. Notably, a linear detection of CMPTX+ EVs was detected in EV+ CM across increasing sample volumes. In contrast the number of CMPTX+ remained stable across increasing volumes of EV- CM. Data represented as Mean ± SEM (*p<0.05; n=3). Significance of analyses were performed by paired-student’s t-test and were assessed exclusively between matched volumes of EV- or EV+ CM.

**Supplemental Figure 5.**
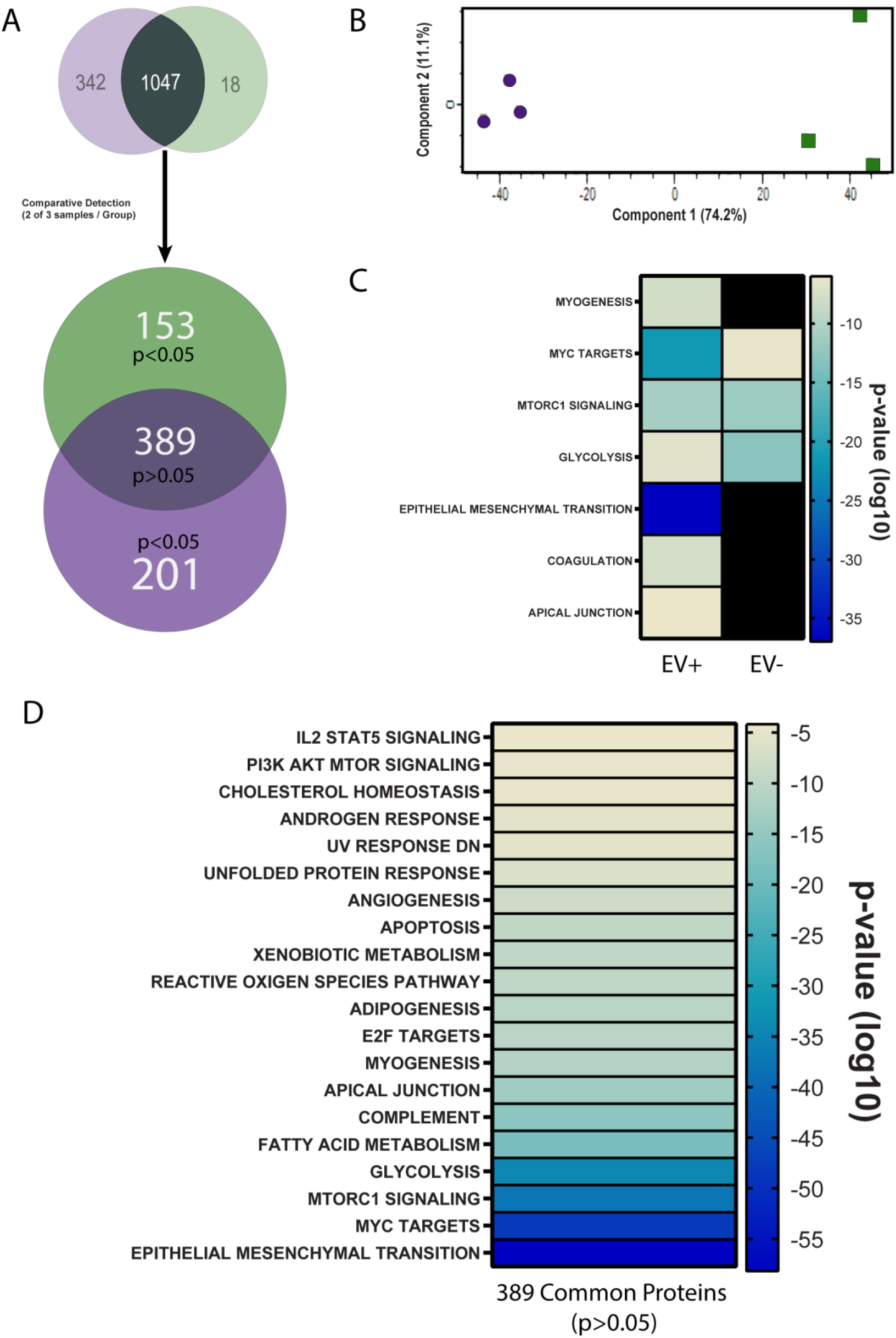
EV+ CM is significantly associated with signaling pathways underlying an epithelial to mesenchymal transition. (A) Of the 1047 proteins identified in EV- or EV+ CM (≥2 of 3 samples), 153 and 201 were significantly enriched (≥2-fold) within EV- and EV+ CM, respectively. 763 proteins were comparably detected in EV- and EV+ CM. (B) PCA analysis demonstrates a distinct proteomic composition of EV- vs EV+ CM. For example, (C) proteins enriched within EV+ CM were significantly associated with epithelial to mesenchymal transitions, referenced against Hallmark Gene Set annotation database. Black boxes in (C) represent annotations which were not significant to either EV- or EV+ CM. Interestingly, (D) detected at comparable levels in EV+ versus EV- CM were also associated with epithelial to mesenchymal transitions.

