## Supplemental Table 1 Antibody List for "Ultrafiltration Segregates Tissue Regenerative Stimuli Harboured Within and Independent of Extracellular Vesicles"

**Supplemental Table 1.** List of primary, secondary and isotype antibodies within manuscript.

| Antibody | Host Species | Antibody Dilution  (Cells) | Company | ID |
| --- | --- | --- | --- | --- |
| CD49f-APC | Mouse | 1:250 | BioLegend | 313615 |
| CD63-Pacific Blue | Mouse | 1:250 | BioLegend | 353011 |
| CD81-Pacific Blue | Mouse | 1:250 | BioLegend | 349515 |
| CD90-APC | Mouse | 1:250 | BioLegend | 328113 |
| APC Isotype | Mouse | 1:250 | BioLegend | 400121 |
| Pacific Blue Isotype | Mouse | 1:250 | BioLegend | 400131 |
