## Supplemental Table 2 Mass Spec Settings for "Ultrafiltration Segregates Tissue Regenerative Stimuli Harboured Within and Independent of Extracellular Vesicles"

**Supplemental Table 2. Mass Spectrometry Parameters.**

| **Parameters** | **Q Exactive** |
| --- | --- |
| **Mass Range (m/z)** | **400-1450** |
| **Isolation Window (m/z)** | **3.0** |
| **MS Resolution** | **35K @ 200 m/z** |
| **MSMS Resolution** | **17.5K** |
| **MS Injection Time (ms)** | **250** |
| **AGC Target (MS)** | **1E6** |
| **AGC (Target MSn)** | **2E5** |
| **Preview Scan** | **n/a** |
| **Threshold (counts)** | **50K** |
| **Underfill Ratio** | **3%** |
| **Data Dependent Acquisition** | **Top 15** |
| **Dynamic Exclusion (s)** | **30** |
| **Exclusion Mass Width (m/z)** | **n/a** |
| **Exclude Isotopes/Monoisotopic**  **Precursor Selection** | **Enabled** |
| **Fragmentation Type** | **HCD** |
| **Normalized Collison Energy** | **23** |
| **Lock Mass (445.120025m/z)** | **Best** |
| **Charge State Rejection** | **Unassigned and +1** |
| **Default Charge State** | **+2** |

(m/z) = mass/charge
